# The influence of HLA genetic variation on plasma protein expression

**DOI:** 10.1101/2023.07.24.550394

**Authors:** Chirag Krishna, Joshua Chiou, Isac Lee, Hye In Kim, Melis Atalar Aksit, Saori Sakaue, David Von Schack, Soumya Raychaudhuri, Daniel Ziemek, Xinli Hu

## Abstract

Polymorphism in the human leukocyte antigen class I (HLA-I) and class II (HLA-II) genes is strongly implicated in susceptibility to immune-mediated diseases. However, the molecular effects of HLA genetic variation, including and beyond antigen presentation, remain unclear. Here we examined the effect of HLA genetic variation on the expression of 2940 plasma proteins using imputed HLA variants in 45,330 Europeans in the UK Biobank. We detected 504 proteins (17.1% of all proteins tested) affected by HLA genetic variation (HLA-pQTL), including widespread *trans* regulation of protein expression by autoimmune disease risk alleles. HLA-pQTL were enriched in gene families related to antigen presentation (e.g. B2M), T cell fate (CD8A; CD4), chemokines (CCL19; CCL21), and NK and macrophage receptors (KIR; LILRA/B), suggesting that HLA polymorphism affects both adaptive and innate immunity. HLA-pQTL also affected expression of diverse proteins with unclear roles in the immune response (e.g. SFTPD, LRPAP1, ENPP6, NPTX1), as well as drug targets for immune-mediated diseases, suggesting complex regulatory roles of the HLA loci. Among *trans* HLA-pQTL, HLA variants explained 0.1-42.9% of the protein expression variance. Fine-mapping revealed that most HLA-pQTL implicated amino acid positions within the peptide binding groove, suggesting that *trans* regulation of plasma protein expression by the HLA loci is primarily a consequence of antigen presentation. We also show that HLA-I and II uniquely affect different proteins and biological mechanisms. Altogether, our data reveal the effects of HLA genetic variation on protein expression and aid the interpretation of associations between HLA alleles and immune-mediated diseases.

## Main

Major histocompatibility complex (MHC) molecules, encoded by the highly polymorphic human leukocyte antigens (HLA) genes in humans, present self and foreign antigens for recognition by T cells and thus comprise the foundation of the adaptive immune response for a wide range of diseases. MHC presentation of viral peptides and neoantigens facilitates clearance of infectious pathogens and cancerous cells, respectively^1–3^; in contrast, aberrant presentation of self-peptides by HLA risk alleles is thought to underly the pathogenesis of autoimmune disease^4, 5^. Polymorphism in the HLA-I (HLA-A, B, and C) and HLA-II (HLA-DRB1, DQB1, DQA1, DPB1, and DPA1) loci is a hallmark of genetic risk for autoimmune diseases; for example, genetic variation in the HLA-II loci explains up to 30% of the genetic heritability in rheumatoid arthritis and type 1 diabetes^4–8^.

Genetic studies linking HLA genes to disease risk have historically been complicated by extreme polymorphism and linkage disequilibrium (LD) in the HLA region. Advances in HLA imputation and statistical association methods have largely overcome these challenges^9, 10^, allowing HLA fine-mapping studies which have pinpointed amino acid positions within the MHC peptide binding groove^5, 7^. These studies lend support to the idea that variation in antigen presentation is likely the pathogenic mechanism by which HLA polymorphism affects disease risk.

Despite much evidence linking HLA to disease risk, the extent to which HLA genetic variation—in particular, the 4-digit alleles that encode the amino acid sequence of the MHC (e.g. HLA DRB1*04:01) —affects quantitative traits such as protein expression levels is incompletely understood^11–14^. Plasma proteins can serve as biomarkers for disease risk and clinical outcomes; and protein quantitative trait loci (pQTL) mapping has improved our understanding of the mechanisms underlying genetic associations with disease^15^. From an immunological perspective, it is conceivable that T cell receptor (TCR) recognition of peptides presented by MHC molecules may lead to changes in protein expression in the corresponding T cell or antigen presenting cell. Moreover, it is possible that certain HLA alleles modulate the strength of the immune response to disease, which in turn may drive differences in protein expression. Thus, a systematic investigation of how HLA variants (e.g. 4-digit alleles and amino acids) affect protein expression would reveal the intermediate molecular effects of HLA polymorphism, and aid the interpretation of associations between HLA polymorphism and disease.

To address this question, we took advantage of data from the UK Biobank, a massive prospective cohort that has facilitated critical insights into the genetic determinants of disease through genome-wide genotyping and health phenotype collection on all participants^16^. A recent flagship pQTL effort led by the UK Biobank Pharma Proteomics Project (UKB-PPP) quantified and performed rigorous quality control of 2940 plasma proteins levels measured with the Olink platform across 54,306 UK Biobank participants^15, 17^. We used these data as input to our study.

To probe the effects of HLA genetic variation on plasma protein expression, we first split the UKB-PPP participants into discovery (N = 34490 individuals) and replication (N = 10840) cohorts, as also performed by the flagship UKB-PPP study^15^ (Methods) (Fig. 1a). In brief, the UKB-PPP discovery cohort is a ‘randomized baseline’ European cohort highly representative of the overall UK Biobank cohort and is neither enriched nor depleted for any particular disease.

**Fig. 1:**
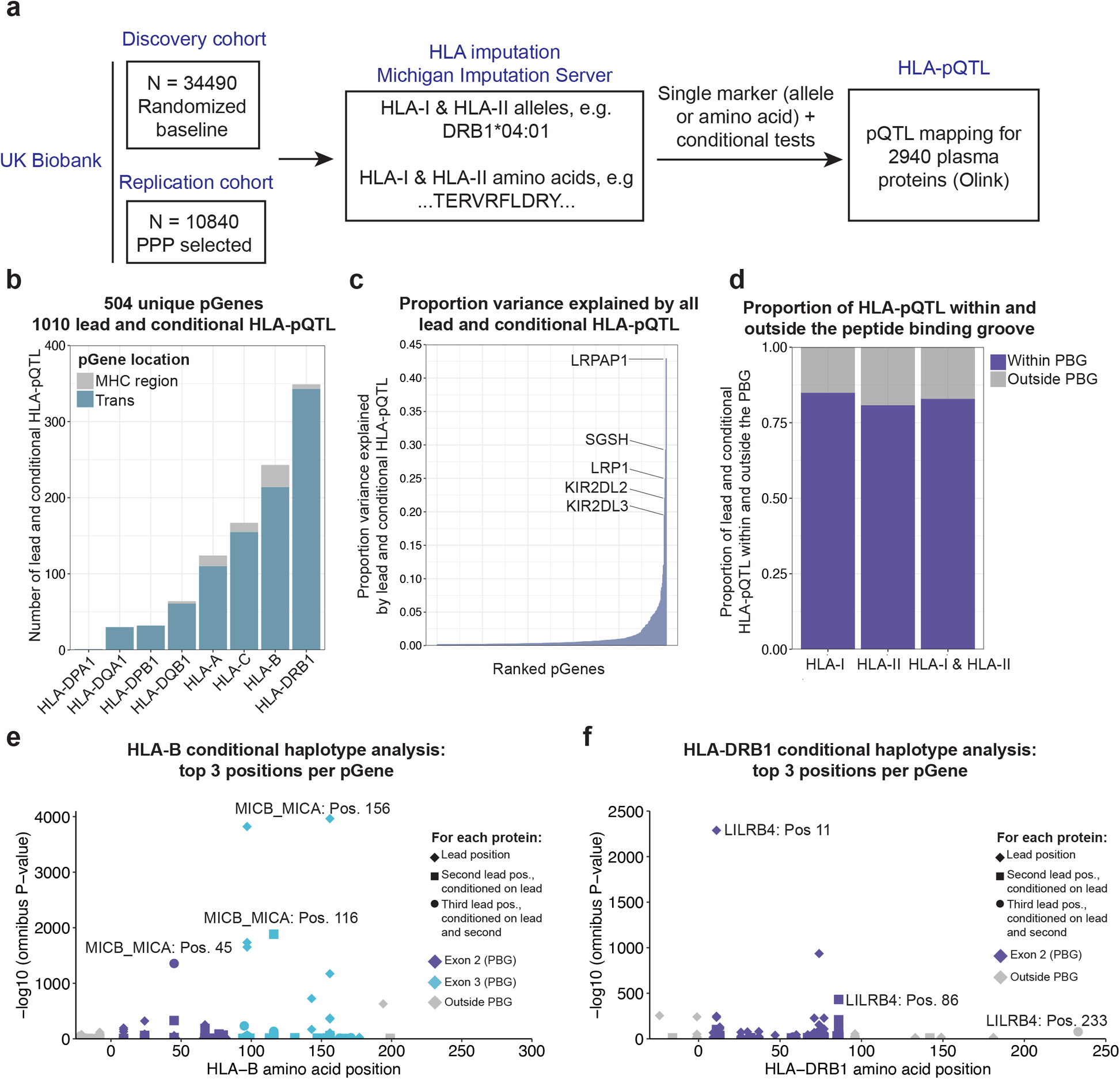
HLA-pQTL in the UK Biobank. **a**, Schematic of the study. We used imputation of 2 and 4-digit HLA alleles and amino acid polymorphisms across 45,330 Europeans from the UKB Pharma Proteomics Project (UKB-PPP) together with plasma protein levels quantified using the OLINK platform as input to pQTL mapping for 2940 proteins. For each protein, we tested each HLA variant (allele or amino acid) independently in a multivariable linear model incorporating covariates as specified by the UKB-PPP. **b**, Number and location of proteins (pGenes) in the discovery cohort affected by lead HLA variants corresponding to each HLA-I and HLA-II locus. **c**, For each ranked pGene in the discovery cohort, proportion of protein expression variance explained by all lead and conditional HLA-pQTL. **d**, Proportion of all fine-mapped lead and conditional HLA-pQTL in the discovery cohort within and outside the peptide binding groove (exons 2 and 3 for HLA-I; exon 2 for HLA-II). **e**, Top three significant (P < 5.0 x 10^−8^) amino acid positions identified via conditional haplotype analysis for HLA-B. The protein with lowest P-value across all proteins tested is labeled. **f**, Top three significant (P < 5.0 x 10^−8^) amino acid positions identified via conditional haplotype analysis for HLA-DRB1. The protein with lowest P-value across all proteins tested is labeled together with its top 3 significant conditional positions.

Following exclusion of individuals for whom genotypes did not pass stringent quality control filters recently established for HLA association studies^10^ (N = 674) (Methods), we used the multi-ancestry reference panel^12, 13^ included in the Michigan Imputation Server to impute HLA 2- and 4-digit alleles and amino acids at MAF ≥ 1% and imputation dosage R^2^ ≥ 0.7 (2284 2-digit alleles, 4-digit alleles, and amino acids total) across 45,330 Europeans in the UK Biobank (‘UKB-PPP cohort’). The imputed variants included 2-digit alleles (e.g. DRB1*04), 4-digit alleles (e.g. DRB1*04:01), and amino acid residues (e.g. AA_DRB1_position13_exon2_FLS) in the classical HLA I and II genes.

To fine map HLA-pQTL, we first tested the association between each imputed HLA variant and covariate-residualized inverse rank-normalized protein expression (Methods) using linear regression models. We included demographic and technical covariates (Methods); additionally, to account for latent sources of variation in the data and the presence of diseases in each cohort, we included the top 20 protein expression PCs as covariates (Extended Data Fig. 1). We set a stringent Bonferroni significance threshold of P ≤ 1.70 x 10^−11^ (5 x 10^−8^ / 2940 proteins tested). To ensure the robustness of lead HLA-pQTL, we performed a permutation analysis in which we randomized protein expression values across individuals in the discovery cohort (Methods). This analysis revealed that our statistical tests had little chance of inflation, as none of the pGenes achieved a p-value in the random data more significant than those observed from the real data (Extended Data Fig. 2).

We identified 504 unique plasma proteins (‘HLA-pGenes’; 17.1% of the 2940 proteins tested, Supplementary Table 1) in the discovery cohort whose expression was affected by at least one HLA variant. To identify independent HLA variants affecting protein expression, we performed conditional analysis by adjusting for the lead HLA variant for each protein, which identified 506 additional independent HLA variants (Supplementary Table 2). Notably, we found that pGenes tagged by HLA-pQTL were less correlated with each other compared to pairs of all other proteins (P = 1.93 x 10^−86^, Extended Data Fig. 3a,b), suggesting that HLA-pQTL do not simply tag correlated proteins. Of the 504 pGenes, 240 had secondary independent associations or more (median 2; range 1-9) (Extended Data Fig. 3c). Strikingly, the majority (480 / 504; 95.2%) of the HLA-pGenes were in *trans* (i.e. proteins outside the MHC region on chromosome 6) (Fig. 1b), suggesting that HLA genetic variation has widespread effects on protein expression. Notably, HLA-DRB1 variants were associated with the largest number of proteins. Prior studies have shown that alleles and fine-mapped amino acid positions in HLA-DRB1 drive risk of autoimmune diseases^5, 7^ and affect the TCR repertoire^18^. Our results corroborate the important role of genetic variation in HLA-DRB1 in influencing both disease risk and intermediate molecular traits.

For each of the 504 pGenes, we quantified the proportion of protein expression variance explained by all independent HLA-pQTL (Fig. 1c). Notably, for some proteins, HLA-pQTL collectively explained more than 20% of the protein expression variance (e.g. 42.89% attributed to LRPAP1; SGSH 29.24%; LRP1 24.90%; KIR2DL2 22.07%). Comparing the variance explained by HLA-pQTL to genome-wide pQTL from the flagship UKB-pQTL study, we found that for several pGenes, most of the phenotypic variance explained by *trans* genetic variation is accounted for by HLA genetic variation (Extended Data Fig. 3d). Next, we quantified the proportion of HLA amino acid variants from our single-marker tests that are located within the peptide binding groove (exons 2 & 3 for the HLA-I loci; exon 2 for the HLA-II loci)^19^. We found that 84.9% of HLA-I pQTL and 80.8% of HLA-II pQTL were located within the peptide binding groove, suggesting that for most proteins, genetic variation in antigen presentation is the likely mechanism by which HLA variants affect protein expression (Fig. 1d).

To pinpoint HLA amino acid positions that affect protein expression, we performed omnibus and conditional haplotype tests for all 2940 proteins (Methods, Fig. 1e,f, Extended Data Fig. 4, and Supplementary Table 3). These analyses revealed strong associations between multiple independent amino acid positions—themselves previously associated with risk of autoimmune diseases^5, 7^—with disease relevant proteins. For example, the most significant signal at the DRB1 locus was for the LILRB4 protein, at HLA-DRB1 position 11 (omnibus P = 1.0 x 10^−2288^), which was previously shown to be associated with risk of rheumatoid arthritis^5^. LILRB4 is a myeloid cell receptor broadly associated with tolerogenic antigen presenting cells^20^. Importantly, the LILR family of proteins (multiple of which were affected by HLA genetic variants in our analysis) are thought to bind MHC molecules, suggesting that HLA-pQTL may reflect cell-cell interactions between APCs and myeloid cells expressing LILR proteins. To identify additional independent amino acid positions in the DRB1-DQB1-DQA1 superlocus (Methods) associated with LILRB4 expression, we conditioned on DRB1 position 11, and observed a second independent position at DRB1 position 86 (omnibus P = 1.0 x 10^−432^), and a third independent position still in DRB1 at position 233 (omnibus P = 1.0 x 10^−78^). Separately from LILRB4, HLA-DRB1 positions 67 and 13 were associated with expression of the LILRA2 protein, demonstrating that HLA-pQTL can affect expression of multiple proteins within the same gene family. Similar analyses for other proteins identified conditionally independent amino acid positions previously associated with disease. For example, expression of CPVL, an enzyme with loosely defined roles in antigen processing^21^, was affected by HLA-DRB1 position 74 (P = 1.0 x 10^−936^), position 11 (P = 6.10 x 10^−37^), and position 71 (P = 7.25 x 10^−13^), all previously associated with risk of type 1 diabetes and rheumatoid arthritis. The unexpectedly strong associations between HLA polymorphism and expression of proteins such as LILRB4/A2, CPVL, CD74, B2M, CD8, CD4, and genes of the KIR family suggest that that the strongest effects of HLA genetic variation on protein expression are related to the immune response, including components of the antigen processing and presentation pathway.

To evaluate the replicability of lead HLA-pQTL from the discovery cohort, we performed HLA-pQTL mapping in the UKB-PPP replication cohort. The replication cohort consists of individuals from the UK Biobank enriched for diseases manually selected by the consortium partners as well as COVID-19 imaging participants, and is thus more heterogeneous in terms of demographics and disease representation than the discovery cohort. We note that the discovery cohort in the flagship UKB-PPP analysis consisted entirely of Europeans while the replication cohort was comprised of individuals with mixed ancestries; thus, we limited the replication cohort to Europeans to ensure consistency with the discovery cohort. We observed high replicability of lead HLA-pQTL, both by significance and concordance of effect sizes (96% at P < 0.05 in the replication cohort; Spearman rho = 0.94 between discovery and replication; Extended Data Fig. 3e and Supplementary Table 4), and combined analysis of both the discovery and replication cohorts together yielded an additional 119 pGenes (623 total, Supplementary Table 5).

We next wished to more closely examine the biology of proteins affected by HLA polymorphism, including 4-digit alleles associated with risk of immune-mediated diseases. We hypothesized that our analyses could help understand the unique regulatory effects of the HLA-I versus HLA-II loci. Indeed, it is generally well-established that HLA-I and HLA-II are recognized by CD8+ and CD4+ T cells respectively, and that HLA-II-restricted antigens are qualitatively different compared to HLA-I antigens^22^. However, the extent to which the *trans* regulatory effects of HLA-I and HLA-II genetic variation differ is unclear.

We first assembled two sets of pGenes from the discovery cohort that were exclusively affected by HLA-I (N = 225 pGenes) or HLA-II (N = 279 pGenes) lead variants. To rule out any potential confounding of HLA-II genetic variation influencing HLA-I-specific pQTL or vice versa, we tested the effects of lead HLA-I pQTL(e.g. the HLA-I pGene set) while also adjusting for all 4-digit alleles corresponding to all HLA-II loci, and vice versa. We found that at P < 5 x 10^−8^, 72.8% (203/ 279) of lead HLA-I pQTL remained significant after adjusting for all HLA-II alleles, and 72.9% of lead HLA-II pQTL remained significant after adjusting for all HLA-I alleles. Next, we performed Gene Ontology enrichment analysis in each pGene set using the full set of Olink panel proteins as the background, and constructed *trans* HLA-pQTL networks separately for HLA-I and HLA-II based on the enriched gene families (Fig. 2a-b, Supplementary Tables 6-7). These analyses showed that inhibitory receptors such as proteins of the KIR family and some members of the LILR family (e.g. LILRB1 and LILRB2) were HLA-I-specific, in addition to proteins involved in TNF binding and NK activity (MICB). In contrast, many chemokines (CCL and CXCL proteins) were HLA-II specific, in addition to general signaling and pattern recognition receptors such as TLR1, IL1/12RB2, and IGF1R, among others. The effect of HLA polymorphism on KIR and pattern recognition receptors suggests that HLA polymorphism also affects components of innate immunity, in addition to its traditional role in the adaptive immune response. Notably, both HLA-I-specific and HLA-II-specific pGenes included known drug targets for autoimmune diseases, infectious diseases, and cancer such as LAG3, PDCD1, TNF, and ACE2 affected by HLA-I; IL18, TREM2, VSIG4, IL12RB2, and TLR1 (HLA-II). The presence of well-known inhibitor receptors, molecules related to T and NK function, cytokines, and chemokines further suggests that the strongest effects of HLA genetic variation on protein expression are related to the canonical immune response (Fig. 2). Moreover, for both HLA-I and HLA-II, we observed effects on proteins with less well-defined functions and relevance to the immune response (Fig. 2c-g). Altogether, these data suggest that HLA genetic variation has complex regulatory effects on protein expression.

**Fig. 2:**
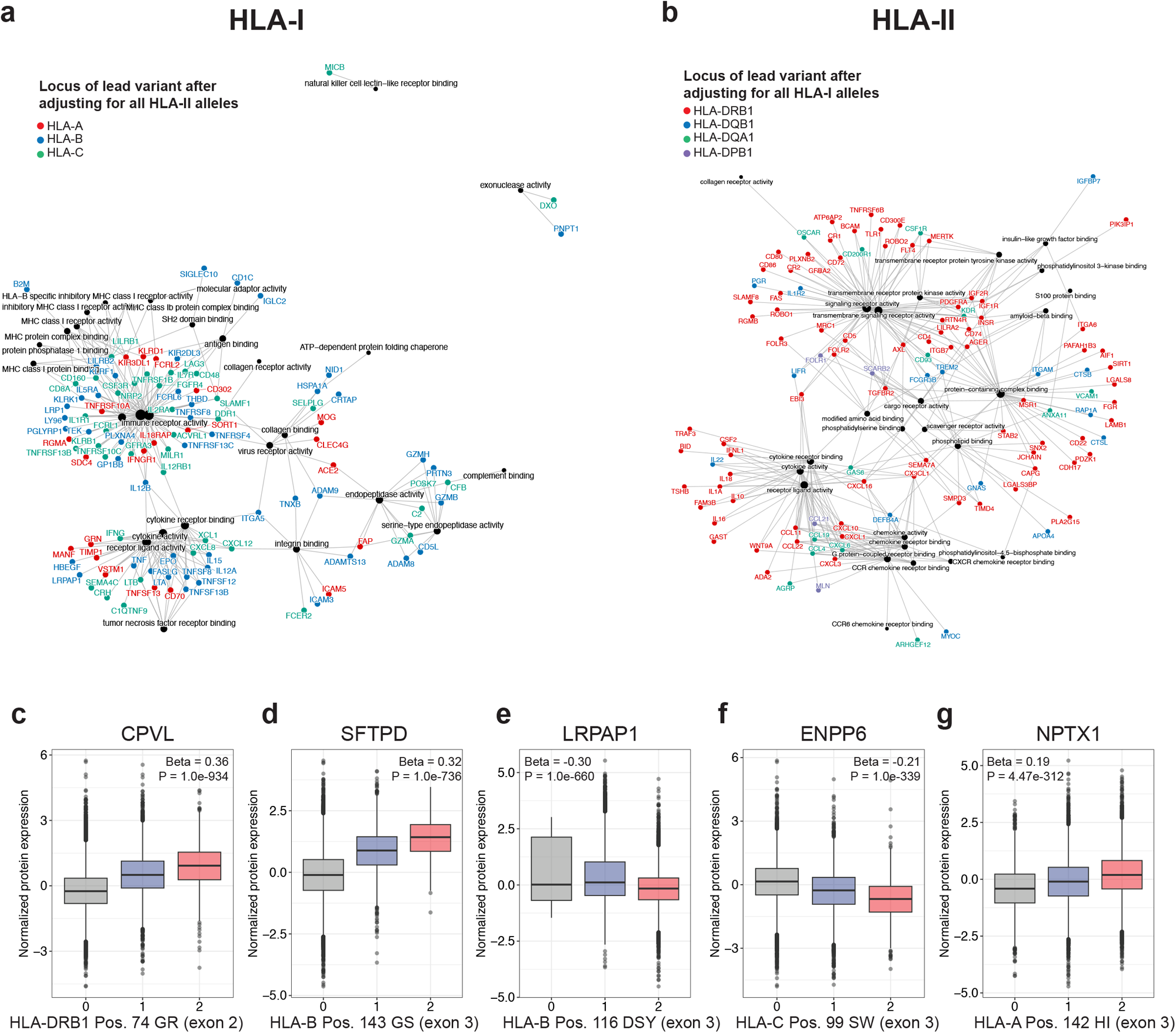
Trans HLA-pQTL networks and gene families affected by HLA-I and HLA-II genetic variation. **a**, Network depicting proteins and gene ontology terms affected by only HLA-I lead variants after adjusting for all HLA-II 4-digit alleles. Edges connect proteins to gene ontology terms. **b**, Network depicting proteins affected by only HLA-II lead variants from after adjusting for all HLA-I 4-digit alleles. Edges connect proteins to gene ontology terms. **c-e**, Selected pGenes from among the top 50 lead HLA-pQTL. These proteins were selected due to their strong associations with HLA genetic variation and unclear or undefined roles in the immune response.

To explore the relevance of HLA-pQTL to human disease, we created an atlas of all proteins affected by HLA-I and HLA-II alleles previously associated with immune-mediated diseases (Table 1 and Supplementary Table 8). Notably, the DRB1*04:01 allele, strongly associated with type 1 diabetes and rheumatoid arthritis, affected expression of multiple proteins and in particular drove higher expression of NFATC3 (β = 0.07; P = 5.49 x 10^−45^), a key transcription factor regulating helper T cell differentiation^23^. The C*06:02 allele, which is a strong genetic determinant of psoriasis, drove higher expression of CD8A (β = 0.21; P = 1.0 x 10^−363^). The strong association between C*06:02 and CD8A expression provide further evidence for the role of CD8 T cells in psoriasis^24^. To further annotate the disease relevance of proteins affected by HLA-pQTL, we asked which pGenes found in our analyses harbor coding variation associated with disease-relevant traits in the UK Biobank, including well-known biomarkers such as BMI, albumin, and urate. We found links between 57 HLA-pGenes and 38 traits (Extended Data Fig. 5). Intriguingly, we detected multiple corneal traits associated with HLA-pGenes, which may support the previously described roles of HLA in corneal immune and epithelial cells^25, 26^.

**Table 1:**
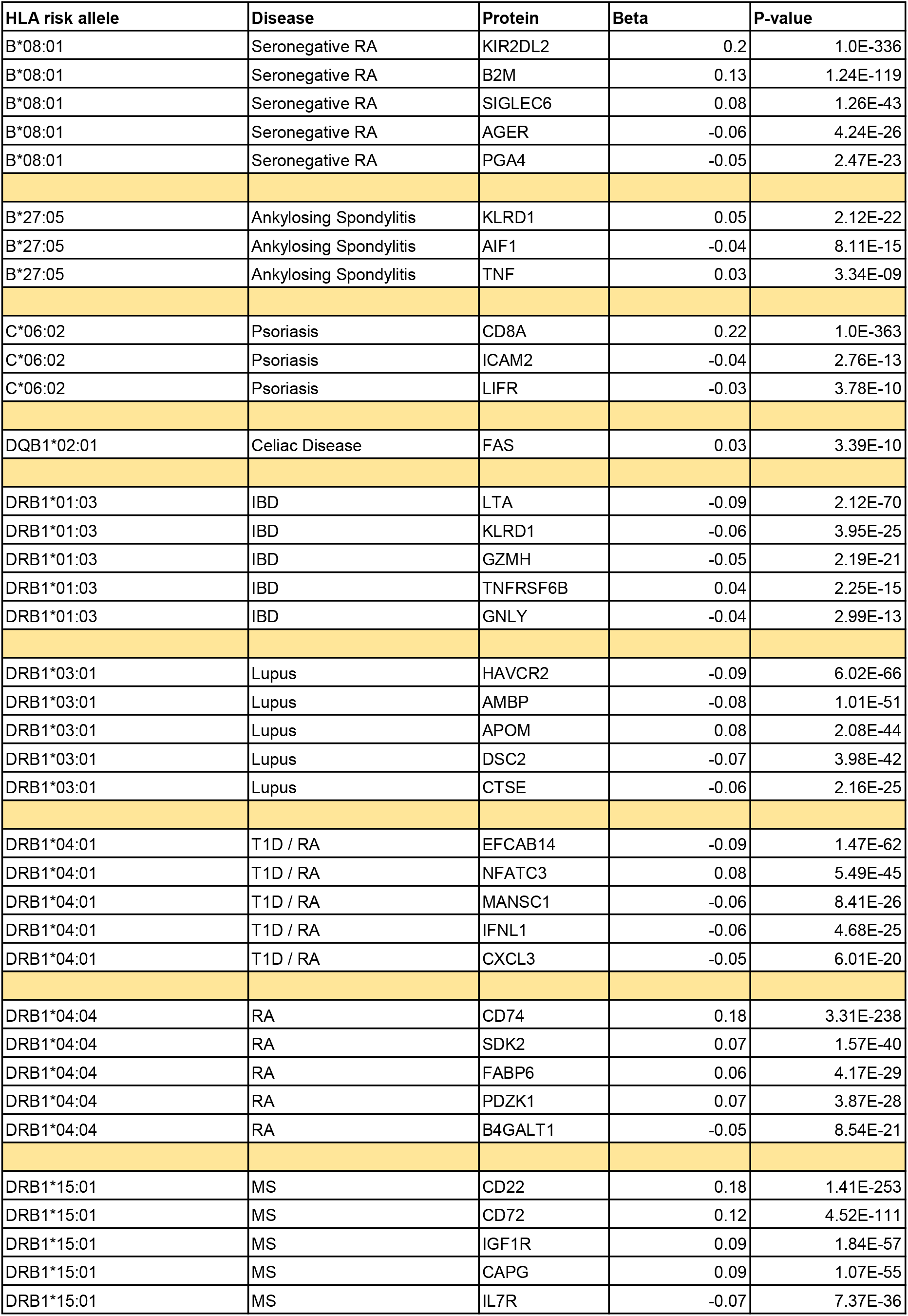
HLA-pQTL involving alleles associated with risk of immune-mediated diseases. The top 5 associations for each allele (when available) are shown. The full conditional HLA-pQTL results for 4-digit alleles only are shown in Supplementary Table 8.

| HLA risk allele | Disease | Protein | Beta | P-value |
| --- | --- | --- | --- | --- |
| B*08:01 | Seronegative RA | KIR2DL2 | 0.2 | 1.0E-336 |
| B*08:01 | Seronegative RA | B2M | 0.13 | 1.24E-119 |
| B*08:01 | Seronegative RA | SIGLEC6 | 0.08 | 1.26E-43 |
| B*08:01 | Seronegative RA | AGER | -0.06 | 4.24E-26 |
| B*08:01 | Seronegative RA | PGA4 | -0.05 | 2.47E-23 |
| B*27:05 | Ankylosing Spondylitis | KLRD1 | 0.05 | 2.12E-22 |
| B*27:05 | Ankylosing Spondylitis | AIF1 | -0.04 | 8.11E-15 |
| B*27:05 | Ankylosing Spondylitis | TNF | 0.03 | 3.34E-09 |
| C*06:02 | Psoriasis | CD8A | 0.22 | 1.0E-363 |
| C*06:02 | Psoriasis | ICAM2 | -0.04 | 2.76E-13 |
| C*06:02 | Psoriasis | LIFR | -0.03 | 3.78E-10 |
| DQB1*02:01 | Celiac Disease | FAS | 0.03 | 3.39E-10 |
| DRB1*01:03 | IBD | LTA | -0.09 | 2.12E-70 |
| DRB1*01:03 | IBD | KLRD1 | -0.06 | 3.95E-25 |
| DRB1*01:03 | IBD | GZMH | -0.05 | 2.19E-21 |
| DRB1*01:03 | IBD | TNFRSF6B | 0.04 | 2.25E-15 |
| DRB1*01:03 | IBD | GNLY | -0.04 | 2.99E-13 |
| DRB1*03:01 | Lupus | HAVCR2 | -0.09 | 6.02E-66 |
| DRB1*03:01 | Lupus | AMBP | -0.08 | 1.01E-51 |
| DRB1*03:01 | Lupus | APOM | 0.08 | 2.08E-44 |
| DRB1*03:01 | Lupus | DSC2 | -0.07 | 3.98E-42 |
| DRB1*03:01 | Lupus | CTSE | -0.06 | 2.16E-25 |
| DRB1*04:01 | T1D / RA | EFCAB14 | -0.09 | 1.47E-62 |
| DRB1*04:01 | T1D / RA | NFATC3 | 0.08 | 5.49E-45 |
| DRB1*04:01 | T1D / RA | MANSC1 | -0.06 | 8.41E-26 |
| DRB1*04:01 | T1D / RA | IFNL1 | -0.06 | 4.68E-25 |
| DRB1*04:01 | T1D / RA | CXCL3 | -0.05 | 6.01E-20 |
| DRB1*04:04 | RA | CD74 | 0.18 | 3.31E-238 |
| DRB1*04:04 | RA | SDK2 | 0.07 | 1.57E-40 |
| DRB1*04:04 | RA | FABP6 | 0.06 | 4.17E-29 |
| DRB1*04:04 | RA | PDZK1 | 0.07 | 3.87E-28 |
| DRB1*04:04 | RA | B4GALT1 | -0.05 | 8.54E-21 |
| DRB1*15:01 | MS | CD22 | 0.18 | 1.41E-253 |
| DRB1*15:01 | MS | CD72 | 0.12 | 4.52E-111 |
| DRB1*15:01 | MS | IGF1R | 0.09 | 1.84E-57 |
| DRB1*15:01 | MS | CAPG | 0.09 | 1.07E-55 |
| DRB1*15:01 | MS | IL7R | -0.07 | 7.37E-36 |

Altogether, our data suggest that HLA risk alleles exert disease-relevant *trans* effects on plasma protein expression. We hypothesize that there may be two primary mechanisms by which HLA polymorphism affects protein expression. First, as our fine-mapping analyses strongly implicate amino acid positions within the peptide binding groove, we suspect that variation in antigen presentation affects T and NK cell recognition of peptide-MHC, which in turn may modulate the expression levels of proteins related to T and NK effector function. Indeed, the expression of a wide array and chemokines and cytokines may be related to T and NK cell recognition of peptides presented by antigen presenting cells^27^. The second potential mechanism by which such changes in protein expression may arise is through cell intrinsic effects in macrophages and other antigen presenting cells (APCs) that demonstrate high expression of HLA. Indeed, this could explain the discovery of proteins involved in antigen processing and presentation, macrophage activating and inhibitory receptors. Future functional studies clarifying the exact mechanisms by which HLA alleles affect protein expression are warranted, especially since several of the proteins affected by HLA genetic variation do not have an immediate connection to T cell or APC biology.

An important caveat of the current study is that only soluble proteins are measured, which may not reflect the fraction of proteins in physiologically relevant compartments. Prior studies have indicated that the soluble forms of at least a few of the proteins included in the Olink panel (e.g. PD1, LAG3) are cleaved directly from the cell surface through membrane shedding^28^. However, where possible, future studies should investigate the effects of HLA genetic variation on membrane-bound proteins.

Moreover, since application of genetic colocalization and Mendelian randomization may not be nuanced enough for the MHC^11^ given the high degree of linkage disequilibrium, mechanistic functional studies may be necessary to validate the extent to which HLA-pQTL are truly causal for disease risk. We emphasize the value of expanding HLA-pQTL analyses to specific cell types, tissues, and diseases pending data availability, as current efforts are limited to blood and a limited panel of proteins. More generally, we hope that our current study will pave the way for investigation of HLA regulatory effects on quantitative traits beyond protein expression.

## Methods

### Study population

The UK Biobank is a prospective, population-scale cohort containing roughly 500,000 participants between the ages of 40-69 at enrollment. All individuals are assessed for demographic, clinical, and phenotypic variables at least once during a baseline visit; genome-wide imputed genetic data are available for all participants. For all analyses presented here, we used individuals from the UK Biobank that were included in the flagship UKB-PPP paper, which contains extensive details on participant selection and characteristics^15^. In brief, the UKB-PPP participants were split into a ‘randomized baseline’ cohort of predominantly healthy individuals; all individuals were of European ancestry. The replication cohort consists of individuals manually selected by the participating pharmaceutical partners enriched for particular diseases; individuals were of multiple ancestries in the replication cohort. To enable replicability of HLA-pQTL across the discovery and replication cohorts and to confidently impute HLA variants across all individuals, we limited the replication cohort to individuals of European ancestry. In total, we analyzed 45,330 individuals from the UKB-PPP, comprised of 34,490 individuals and 10,840 individuals in the discovery and replication cohorts, respectively.

### Plasma protein measurements and quality control

Full details on the measurement, processing, and quality control of Olink proteomic data for all UKB-PPP participants can be found in the supplemental information of the flagship UKB-PPP analysis. We followed the quality control steps on the protein expression data as the UKB-PPP flagship paper. The Olink technology is an antibody-based assay, with 2940 proteins measured across 8 panels (Cardiometabolic, Cardiometabolic II, Inflammation, Inflammation II, Neurology, Neurology II, Oncology, Oncology II), each focused on specific proteins selected by the UKB-PPP consortium partners across broad disease areas. Within each of the eight panels, the flagship study used principal component analysis (PCA) to remove individuals with PC1, PC2, or NPX (protein expression value log_2_ normalized relative to Olink assay controls) greater than 5 standard deviations from the mean. The flagship study confirmed that there were no abnormalities in protein coefficients of variation (CVs); the percentage of variability attributable to batch and plate effects was also negligible. For each protein, we performed inverse rank-normalized transformation, regressed all covariates, and scaled the protein expression values to mean 0 and standard deviation 1 prior to pQTL mapping.

We used the inverse rank-normalized protein levels subject to all processing and quality control measures used in the flagship study for pQTL testing in our study. The batch and plate of each sample and protein were recorded and used as input to pQTL modeling (see section below, ‘Linear models for mapping HLA-pQTL’). Furthermore, as performed in the flagship study, we subtracted REGENIE LOCO values from the residualized protein expression values to account for local polygenic effects and sample relatedness^29^.

### Quality control of SNP genotypes from the UK Biobank and HLA imputation

We followed recently published guidelines^10^ (https://github.com/immunogenomics/HLA_analyses_tutorial/blob/main/tutorial_HLAQCImputation.ipynb) for genotype QC and HLA imputation using the genome-wide imputed SNP array data from the UK Biobank as input. We used PLINK to remove duplicated SNPs and used snpflip to correct reverse/forward strand flips. We then performed a first round of SNP QC to remove poor quality variants, i.e. those with high genotype call failure rate and violation of Hardy-Weinberg equilibrium (PLINK parameters –geno 0.1 – hwe 1e-10). Next, we filtered samples with greater than 3% missingness (--mind 0.03), based on examining the distribution of genotype missingness across the full cohort. We then removed individuals with genome-wide heterozygosity three standard deviations above the mean; samples with too much or too little heterozygosity may reflect contaminated DNA samples or inbreeding, respectively. We next removed individuals with high genetic relatedness (IBD PI_HAT > 0.9) and performed genome-wide identity-by-state (IBS) computation after removal of the MHC region (chr6: 28000000-34000000). In the next round of SNP QC, we first examined the distribution of SNP missingness and removed variants with more than 3% missingness (--geno 0.03). We then confirmed the accuracy of our quality control measures by comparing allele frequencies between the UK Biobank and 1000 genomes, and confirmed that allele frequencies were highly correlated both genome-wide and within the MHC region (P < 2.2e-16, Spearman rho 0.92 for both). We then removed any SNPs with more than 20% difference in allele frequency between the UK Biobank and 1000 genomes. In total, 7391 variants in the MHC passed QC and were used as input to HLA imputation. We performed HLA genotype imputation with Minimac4 as implemented in the Michigan Imputation Server, using the QC’d MHC SNP data from the UK Biobank as input. We used the most recent reference panel (‘Four-digit Multi-ethnic HLA reference panel v2) for imputation^10, 13^. After imputation, we retained 2284 HLA variants at MAF ≥ 1% and imputation dosage R^2^ ≥ 0.7, comprised of 74 2-digit alleles, 93 4-digit alleles, and 2117 amino acid polymorphisms.

### Covariates and linear models for HLA-pQTL mapping

For each of the 2940 proteins, we fit a multivariable linear regression model testing the imputed dosage (continuous value between 0 and 2) for each of the 2284 HLA variant independently together with all covariates included in the flagship UKB-PPP study. In the discovery cohort, these covariates were age, age^2^, sex, age*sex, age^2^*sex, batch, UKB centre, UKB genetic array, time between blood sampling and measurement, and the top 20 principal components (PCs) of genetic ancestry. Significance in the discovery cohort for lead HLA-pQTL was defined as P ≤ 1.70 x 10^−11^ (5 x 10^−8^ / 2940 proteins tested). To control for additional sources of variation—including the presence of diseases in the cohort—we performed PCA on the normalized protein expression values across all individuals in the discovery cohort, performing mean imputation for individuals without particular protein values. Across all proteins, the mean proportion of individuals with missing values was 0.032 (median 0.03, I.Q.R. 0.02-0.04). In addition to the covariates included from the flagship UKB-PPP study, we also included the top 20 PCs of protein expression in the linear regression models. We repeated this procedure in the replication cohort. Linear models in the replication cohort included an additional covariate detailing whether the participant was pre-selected by the UKB-PPP consortium members or COVID imaging study. Normalized protein expression values were obtained from the flagship study, in addition to the leave one chromosome out (LOCO) values for each protein. Covariates were regressed from the protein expression values and HLA variant dosages, consistent with the Regenie LOCO scheme.

### Evaluation of diseases in the UK Biobank captured by principal components of protein expression

The flagship UKB-PPP study computed associations between each of the 2940 proteins and the most common diseases (defined by ICD-10 codes) in the UK Biobank. We converted these associations to Z-scores, and performed Spearman correlation of these Z-scores with the protein loadings for the top 20 protein expression PCs.

### Pairwise correlations of protein expression

To evaluate whether HLA-pGenes might simply be proteins that are strongly correlated with one another, we computed pairwise correlations of protein expression across all individuals and all 2940 proteins. We detected weak correlations between most proteins (median Spearman rho 0.05, I.Q.R. 0.02-0.13, Extended Data Fig. 3a).

### Single marker tests

To identify lead and conditional single marker HLA-pQTL, we tested each of the 2284 HLA variants (2-digit alleles, 4-digit alleles, and amino acid polymorphisms) independently for association with protein expression levels using the linear regression models and covariates described above. We adopted a conservative threshold for significance of lead pQTL of P ≤ 1.70 x 10^−11^ determined by dividing the genome-wide significance threshold (P ≤ 5 x 10^−8^) by the total number of proteins tested (N = 2940). For conditional analyses, we conditioned on the lead HLA variant and repeated the linear regression until no variant was significant at P ≤ 5 x 10^−8^. To assess the replicability of lead HLA-pQTL, we performed these same analyses in the replication cohort, and checked the significance (P < 0.05) and direction of effect for all lead HLA-pQTL.

### Permutation analysis

To assess the robustness of HLA-pQTL in the discovery cohort, we performed 1000 permutations per protein randomizing protein expression values across donors on each iteration, and testing all HLA variants in each permutation. We retained the lead variant from each iteration to construct an empirical null distribution for each protein.

### Conditional haplotype analysis

To determine whether groups of amino acids inherited together jointly influence variation in protein expression, we used phased best guess amino acid genotype information to conduct conditional haplotype tests. First, for each protein, we evaluated the association between each amino acid position and protein expression using the linear regression models and covariates described above. This first linear regression model includes all amino acid polymorphisms at the indicated position, created by grouping all 4-digit alleles for the particular locus into *M* groups based on the amino acid at the tested position. To obtain an omnibus P-value for a particular amino acid position, we compared this first linear regression model to a null model without any of the *M* groups, using the anova() function in R. Following identification of the amino acid position with minimal P-value (P < 5.0e-08) from the first round of conditional haplotype analysis, we repeated this analysis sequentially conditioning on additional amino acid positions within the locus of the lead position until no position reached significance (i.e. if the lead position is in HLA-A, sequentially condition on other positions within HLA-A). Given the complexity of the DRB1-DQB1-DQA1 super-locus, for all proteins for which the lead position was within DRB1, DQB1, or DQA1, we tested all amino acid positions in DRB1, DQB1, and DQA1 when adjusting for the lead and additional independent variants, as performed previously^7^. We limited our conditional haplotype analyses to the top three conditionally independent amino acid positions per protein.

### Proportion of protein expression variance explained by HLA-pQTL vs genome-wide trans SNP pQTL

We wished to assess the proportion of trans genetic heritability explained by HLA genetic variation vs all other SNPs. Thus, we first used the anova() function in R to compute the variance explained by all lead and conditional HLA-pQTL, and compared these estimates to the variance explained by all lead and conditional SNP pQTL from the flagship UKB-PPP study.

### HLA-I and HLA-II-specific pGenes and network analyses

For the initial assessment of proteins uniquely affected by HLA-I or HLA-II genetic variation, we assembled two sets of mutually exclusive pGenes—those with a lead variant in HLA-I, and those with a lead variant in HLA-II. To rigorously rule out any confounding of HLA-II genetic variation for HLA-I-specific pGenes and vice versa, in each pGene set, we tested the lead HLA variant adjusting for all 4-digit alleles of the other class (i.e. for a pGene with HLA-I lead pQTL, we included all 4-digit HLA-II alleles as covariates in the linear regression model). pGenes that remained significant (P ≤ 5 x 10^−8^) after this analysis were used as input to Gene Ontology enrichment analyses, using the full Olink panel (N = 2940 proteins) as the background. Significant (FDR P < 0.05) Gene Ontology terms were used as input to the cnetplot() function in the clusterProfiler package.

### Coding variation in HLA-pGenes associated with traits

To explore the association of coding variation in HLA-pGenes with traits, we retrieved data from the genebass browser, and searched for all HLA-pGenes with significant (SKAT-O P < 2.5e-07) associations.

## Supplementary Information

**Supplementary Table 1:** Lead HLA-pQTL from the UKB-PPP discovery cohort

**Supplementary Table 2:** Conditional HLA-pQTL from the discovery cohort

**Supplementary Table 3:** Significant multiallelic amino acid positions from omnibus and conditional haplotype tests in the discovery cohort

**Supplementary Table 4:** Lead HLA-pQTL from the UKB-PPP replication cohort

**Supplementary Table 5:** Lead HLA-pQTL from the combined UKB-PPP discovery and replication cohorts

**Supplementary Table 6:** Gene ontology enrichment results for HLA-I-specific pGenes

**Supplementary Table 7:** Gene ontology enrichment results for HLA-II-specific pGenes

**Supplementary Table 8:** Conditional HLA-pQTL from the discovery cohort based on conditional analysis of 4-digit alleles only (to support Table 1)

## Data availability

Summary statistics for all analyses are available in the supplementary materials.

## Acknowledgements

The authors acknowledge Melissa Miller, Anders Malarstig, and the UKB-PPP for their contributions to the design of the UKB Olink dataset. The authors also acknowledge all members of the Systems Immunology team at Pfizer for their helpful comments. This research has been conducted using the UK Biobank Resource under Application Numbers 65851 and 26041.

---

**Extended Data Fig. 1:**
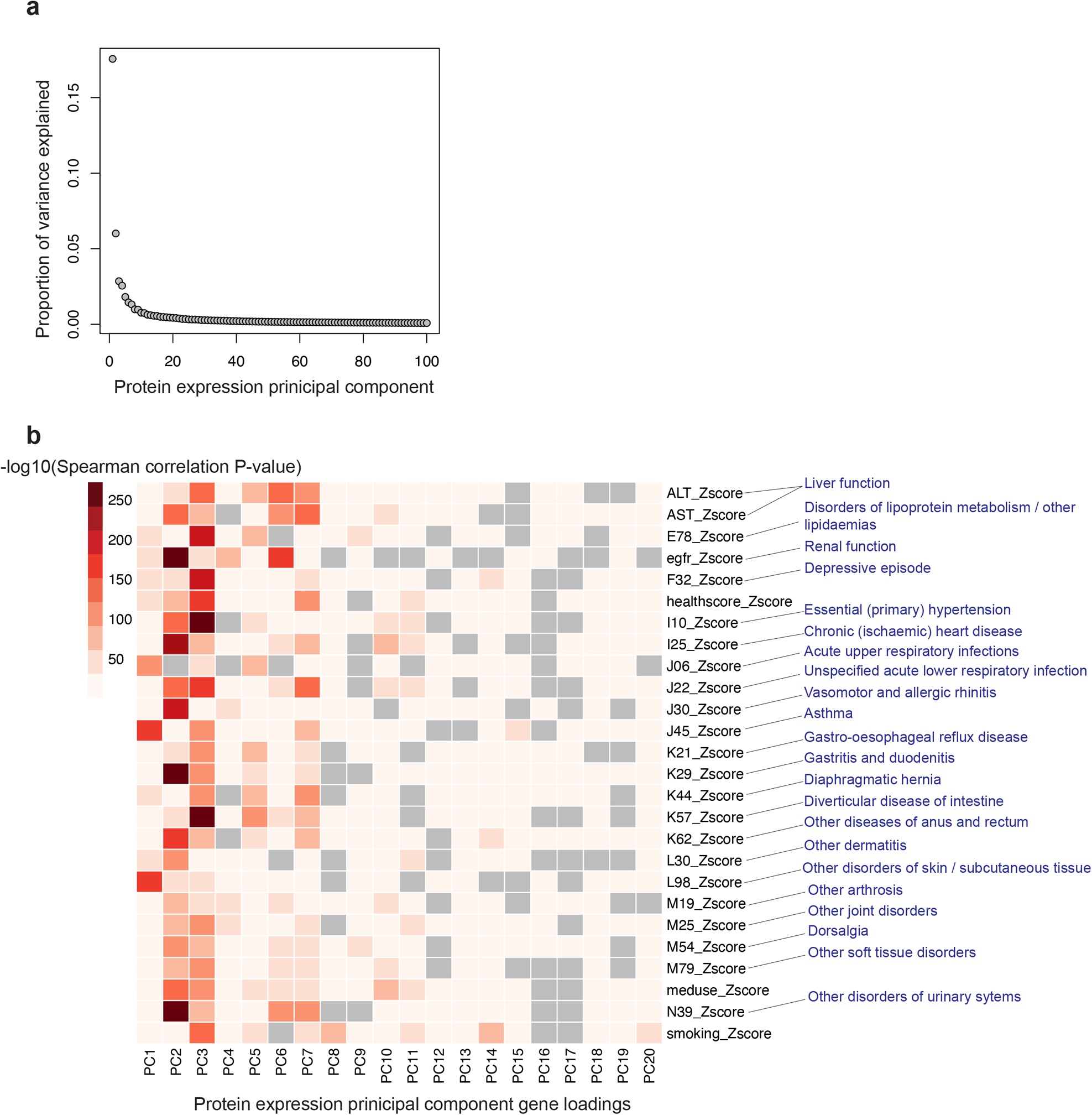
Principal component analysis of plasma protein expression in the UK Biobank. **a**, Proportion variance explained by the top 100 principal components (PCs) of protein expression in the discovery cohort. The top 20 protein expression PCs were used as covariates in the HLA-pQTL linear models. **b**, Spearman correlation of the gene loadings for top 20 protein expression PCs with gene – disease associations (represented as Z-scores) computed by the UKB-PPP, for the most prevalent diseases in the discovery cohort. Gray indicates a non-significant (P-value > 0.05) association. Heatmap rows depict ICD10 code; blue text indicates description of ICD-10 code.

**Extended Data Fig. 2:**
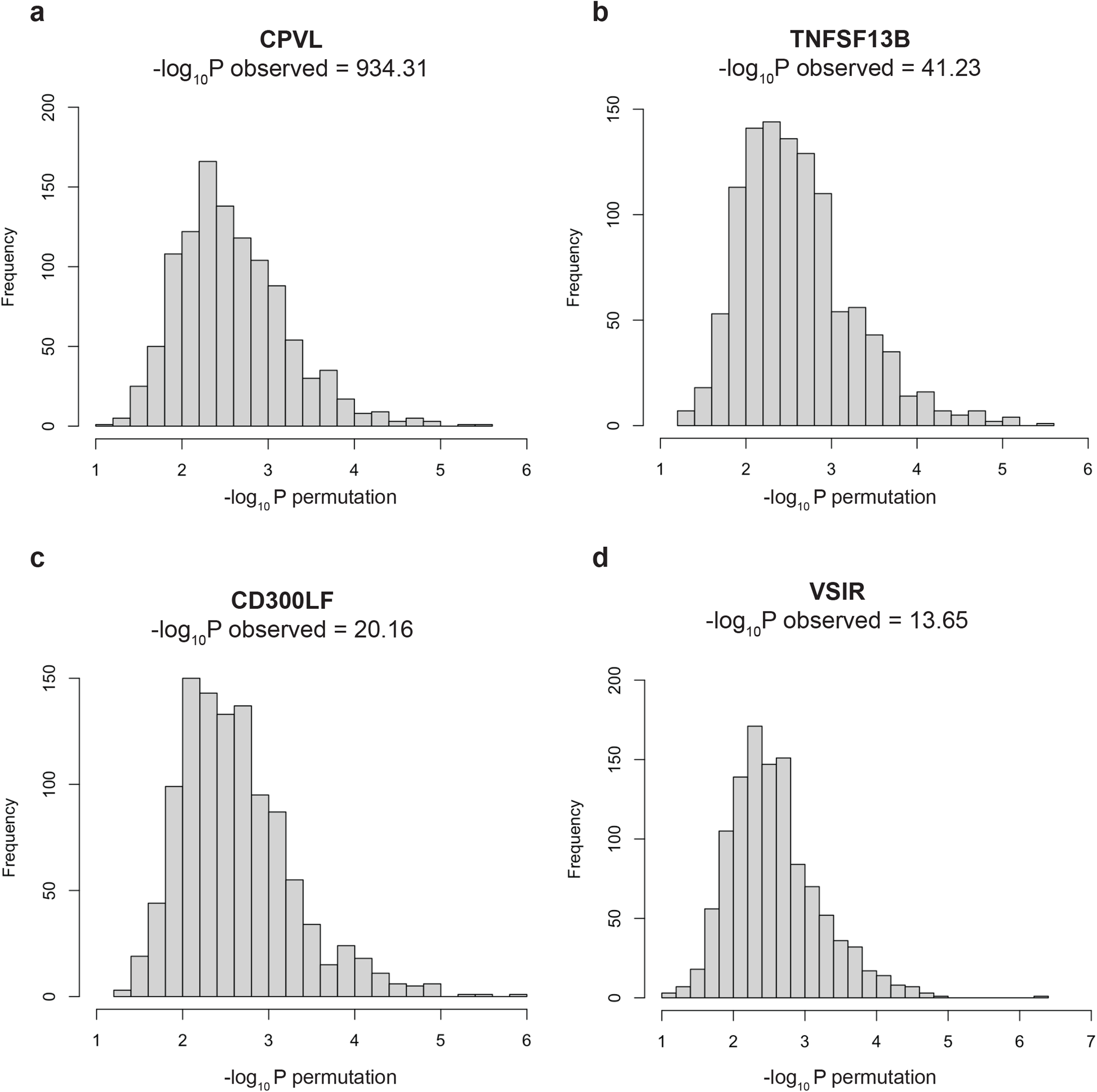
Permutation analysis to assess robustness of lead HLA-pQTL in the discovery cohort. **a-d**, For 4 selected proteins, histograms of −log_10_ permutation p-values in which protein expression values were randomized across participants in the discovery cohort. For each permutation iteration (1000 total for each protein), all HLA variants were tested, and the lead variant (variant with minimal p-value) was retained to construct the null distribution plotted here. For all proteins, no permutation iterations achieved a −log_10_ permutation p-value greater than the observed −log_10_ p-value, listed for each protein.

**Extended Data Fig. 3:**
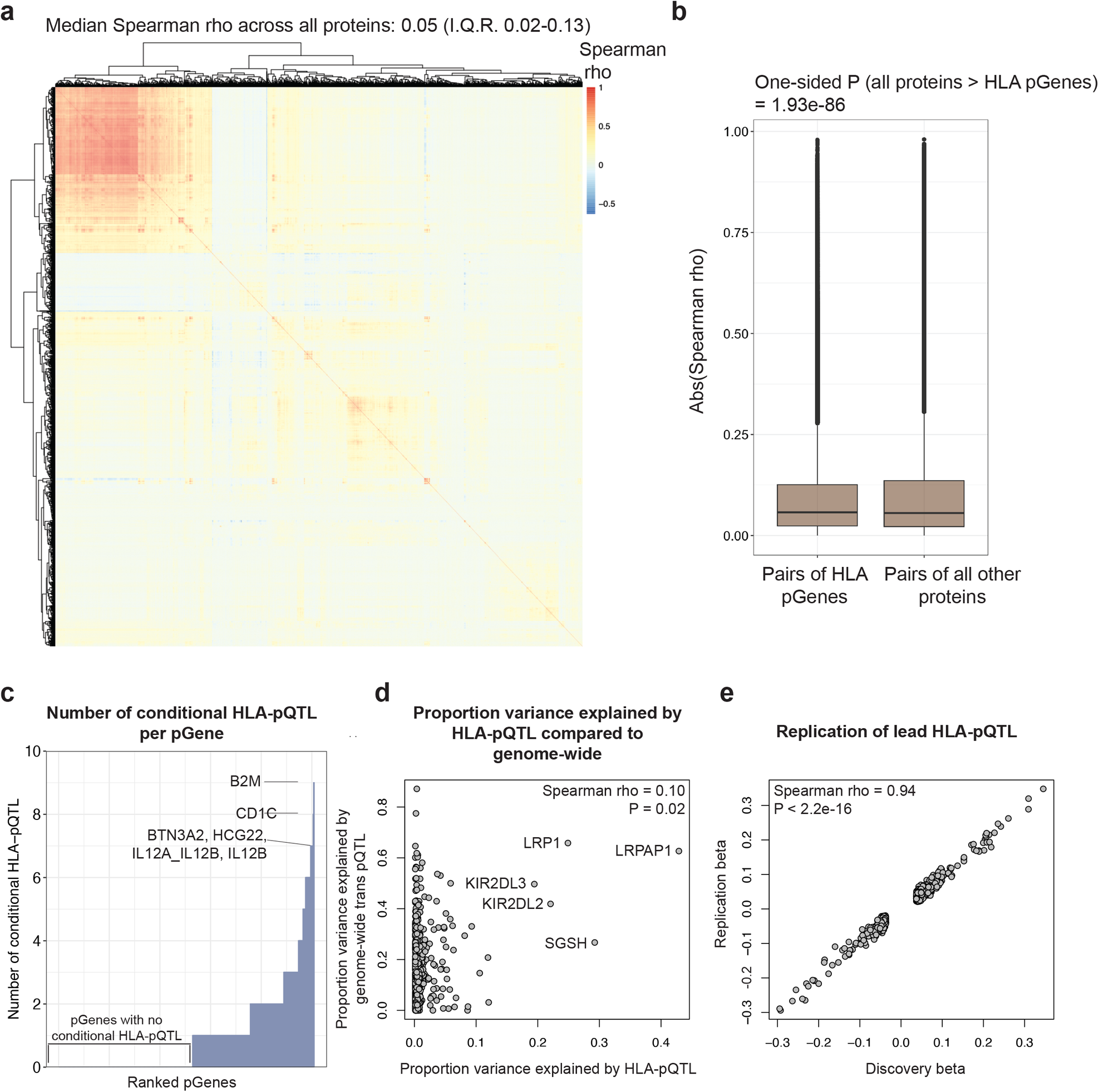
Pairwise protein correlations and additional characteristics of HLA-pQTL. **a**, Pairwise Spearman correlations computed across all 2940 proteins. **b**, Spearman correlations for HLA-pGenes compared to all other pGenes. P-value calculated using one-sided Wilcoxon test. **c**, Number of conditional single-marker HLA-pQTL across all 504 pGenes. **d**, Proportion of protein expression variance explained by lead and conditional HLA-pQTL compared to genome-wide trans SNP pQTL from the flagship UKB-PPP analysis. **e**, Concordance of effect sizes for HLA-pQTL that were nominally significant (P < 0.05) in the replication cohort.

**Extended Data Fig. 4:**
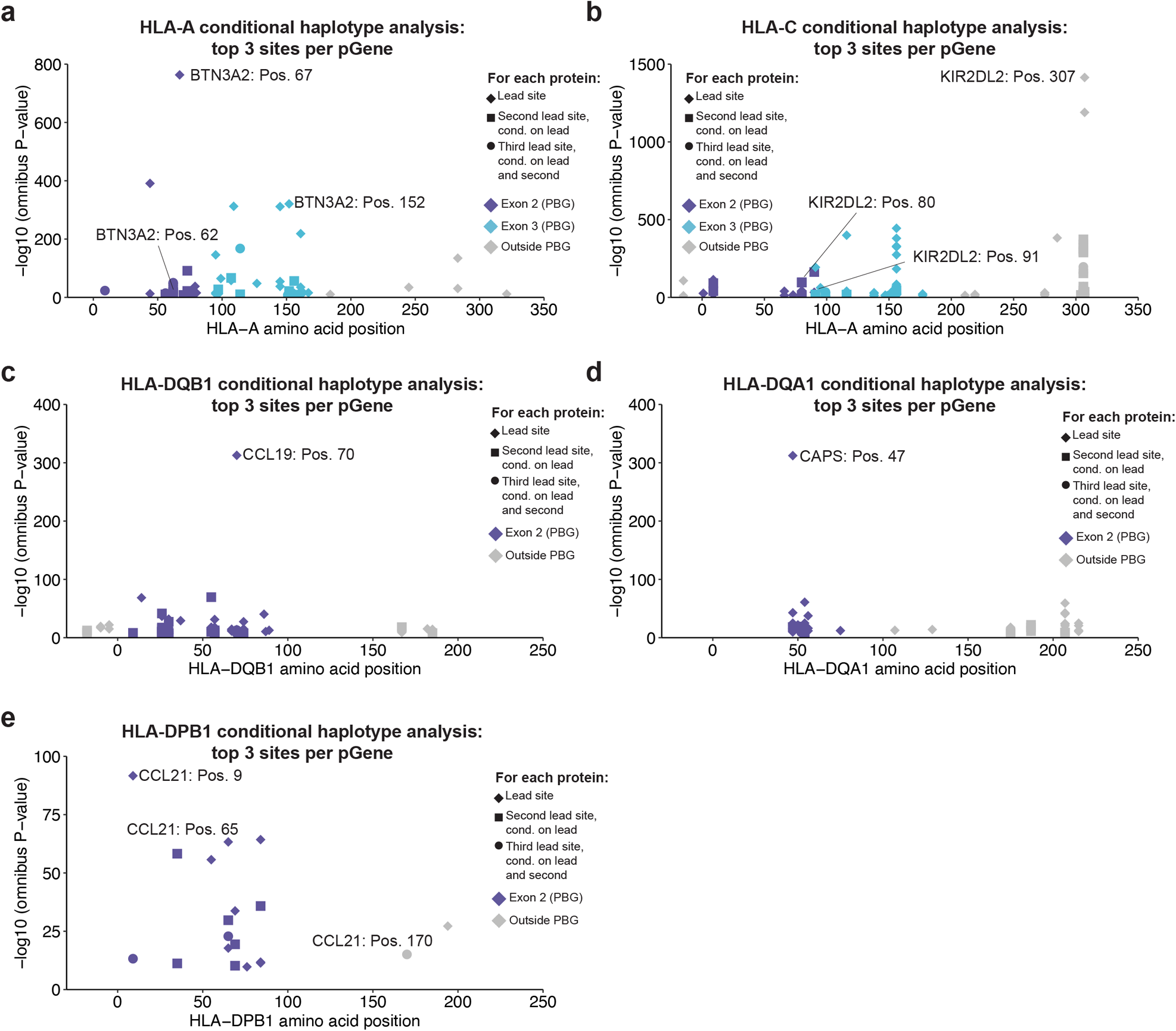
Significant HLA-pQTL identified via conditional haplotype analysis for loci not shown in Fig. 1. **a-e**, Top three significant (P < 5.0 x 10^−8^) amino acid sites identified via conditional haplotype analysis shown for each locus, where available. The protein with lowest P-value across all proteins tested is labeled together with its top 3 significant conditional sites.

**Extended Data Fig. 5:**
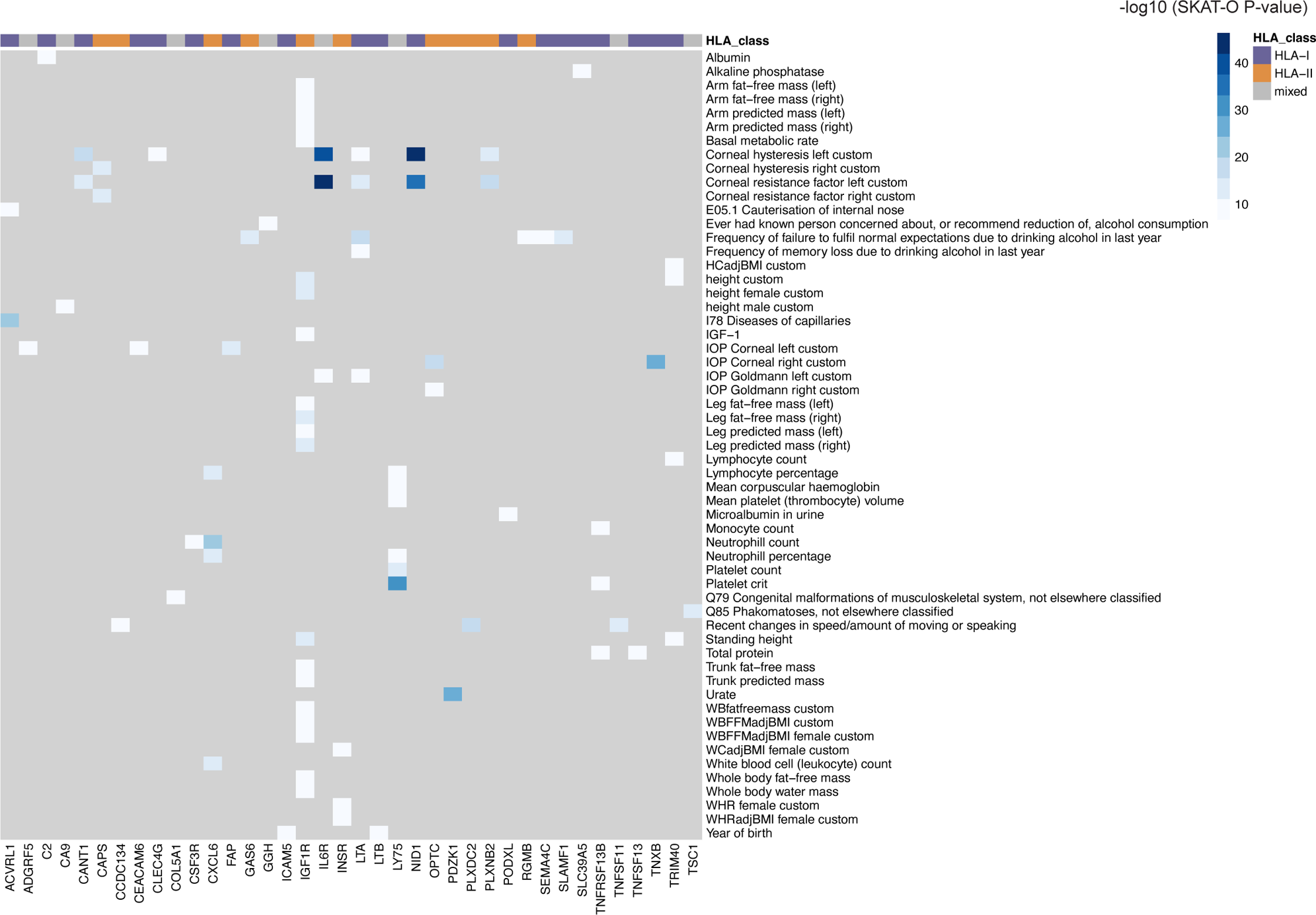
Disease associations for HLA-pGenes obtained via burden tests using genetic variation from exome sequencing. Heatmap shows significant (SKAT-O P < 2.5 x 10^−7^) associations from the genebass browser. Gray indicates non-significant association.

